# Impact of meningioma and glioma on whole-brain dynamics

**DOI:** 10.1101/2025.01.17.633524

**Authors:** Albert Juncá, Anira Escrichs, Ignacio Martín, Gustavo Deco, Gustavo Patow

## Abstract

Brain tumors, particularly meningiomas and gliomas, can profoundly affect neural function, yet their impact on brain dynamics remains incompletely understood. This study investigates alterations in normal brain function among meningioma and glioma patients by assessing dynamical complexity through the Intrinsic Ignition Framework. We analyzed resting-state fMRI data from 34 participants to quantify brain dynamics using intrinsic ignition and metastability metrics. Our results revealed distinct patterns of disruption: glioma patients showed significant reductions in both metrics compared to controls, indicating widespread network disturbances. In contrast, meningioma patients exhibited significant changes predominantly in regions with substantial tumor involvement. Resting-state network analysis demonstrated strong metastability and metastability/ignition correlations between regions in controls, which were slightly weakened in meningioma patients and severely disrupted in glioma patients. These findings highlight the differential impacts of gliomas and meningiomas on brain function, offering insights into their distinct pathophysiological mechanisms. Furthermore, these results show that brain dynamics metrics can be effective biomarkers for identifying disruptions in brain information transmission caused by tumors.

## 1 Background

Glioma and meningioma tumors represent two of the most prevalent types of primary brain tumors, each arising from distinct cellular origins and exhibiting unique clinical characteristics [1, 2, 3]. Gliomas originate from glial cells, integral components of the central nervous system, and play crucial roles in maintaining neuronal function and overall brain health [4]. Gliomas pose significant challenges in neuro-oncology due to their aggressive nature and diverse clinical manifestations. They are known for their infiltrative characteristics, often diffusing into surrounding brain tissue, which complicates treatment efforts [5]. The World Health Organization (WHO) classifies adult gliomas into three primary types based on their cellular origin: astrocytoma, oligodendroglioma, and glioblastoma. These are further categorized from low-grade (I and II) to high-grade (III and IV) based on histological features and aggressiveness [2]. By contrast, meningiomas arise from the meninges, the protective membranes surrounding the brain and spinal cord [6]. The WHO classifies these tumors into three grades based on histological characteristics: Grade I (benign), typically slow-growing with well-defined borders; Grade II (atypical), displaying more aggressive features; and Grade III (anaplastic or malignant), the most aggressive subtype [2]. Both gliomas and meningiomas can disrupt normal neuronal dynamics by various methods such as infiltrating or compressing brain tissue, leading to a wide range of neurological symptoms and impairments [6, 7]. Understanding whole-brain network disruptions and their functional consequences caused by these tumors is crucial for developing effective therapeutic strategies and understanding their impact on psychomotor and cognitive functions.

The human brain is a highly dynamic and complex organ, that orchestrates cognitive and physiological processes through complex networks of neurons [8]. Understanding the patterns of neuronal activity and their interactions over time, referred to as brain dynamics, is crucial for unraveling the mechanisms underlying neurological functions and dysfunctions. These dynamics, observable through neuroimaging techniques such as functional magnetic resonance imaging (fMRI), reveal how brain regions communicate and coordinate, even in resting states [9, 10, 11]. This understanding is vital for addressing neurological and psychiatric disorders, where abnormal neuronal oscillations and disrupted network connectivity are often pivotal, with significant implications for conditions such as epilepsy, Alzheimer’s disease, and schizophrenia [12, 13, 14].

Advances in computational neuroscience have significantly enhanced our ability to analyze and model the brain’s architecture and dynamics in healthy and diseased states [15]. Whole-brain studies present a multifaceted challenge and opportunity, as the complex network of interconnected neurons and glial cells orchestrates a myriad of processes underlying cognition, perception, and behavior [8, 16]. Understanding the brain’s functioning provides essential insights into the fundamental principles governing neural dynamics and information processing. Assessing whole-brain function in patients with neurological disorders, including brain tumors, offers crucial information about the disease on both individual and collective levels. Particularly, studying brain dynamics in patients with brain tumors such as meningiomas and gliomas is fundamental due to the significant disruptions they cause to neural networks, with recent evidence suggesting these effects extend far beyond the immediate tumor location. Aerts et al. [17] examined both computational and empirical studies, finding that brain tumors, especially low-grade gliomas, can cause widespread alterations in functional network topology, affecting network segregation and integration. Hart et al. [18] further demonstrated that glioblastomas produce long-range gradients in cortical function. These tumors alter normal neuronal dynamics and connectivity through various mechanisms, such as infiltrating and compressing brain tissue, which profoundly affects cognitive and psychomotor functions—key aspects of patient quality of life.

While these studies have provided valuable insights, a critical need remains for more comprehensive research to fully understand the complex interactions between brain tumors and neural dynamics. Further investigation is necessary to elucidate the specific mechanisms by which different types of tumors affect brain function across various scales and networks. This knowledge is essential for developing targeted therapeutic strategies to mitigate the adverse effects of tumors on brain function. Additionally, this research can contribute to a broader understanding of how brain tumors interact with and alter neural circuits, providing insights that could apply to other neurological conditions with similar network disruptions.

In this study, we investigated the dynamical complexity across the whole-brain network and within resting-state networks (i.e., visual, somatomotor, dorsal, ventral, limbic, frontoparietal, and default mode network) in patients suffering from glioma and meningioma brain tumors to understand the complex disruptions within brain networks caused by these diseases. Our investigation was structured into three main analyses: (i) an examination of whole-brain dynamics at the subject level to investigate differences between groups, (ii) at the region level to investigate differences in tumor-affected regions on whole-brain dynamics and (iii) across resting-state networks to investigate the relationships between different networks and their disruptions.

## 2 Results

### 2.1 Overview

This study analyzed pre-operative data [19] from 34 participants: 10 control subjects, 10 glioma patients, and 14 meningioma patients recruited at Ghent University Hospital (Belgium) from May 2015 to October 2017, meeting specific inclusion criteria including age ≥ 18 years and diagnosed supratentorial brain tumors (see Table 1). MRI data acquisition included resting-state functional MRI (see Section 4.2). The imaging data were processed using FSL and the Python Nilearn library. Two brain parcellations were employed: the Shen268 atlas (268 regions) [20] for detailed analysis and the Yeo7 atlas [21] (7 resting-state networks) for network-level insights.

**Table 1.** Demographic and Clinical Characteristics of Study Participants.

| Characteristic | Control (n=10) | Glioma (n=10) | Meningioma (n=14) |
| --- | --- | --- | --- |
| Age, years | 59.7 ± 10.1 | 47.7 ± 12.3 | 60.4 ± 12.3 |
| Female sex, n (%) | 3 (30%) | 4 (40%) | 11 (78.6%) |
| Tumor volume, $\text{cm}^3$ | - | 23.43 ± 18.24 | 17.01 ± 29.17 |
| WHO Grade, n (%) |  |  |  |
| I | - | - | 13 (92.9%) |
| II | - | 6 (60%) | 1 (7.1%) |
| III | - | 3 (30%) | - |
| II-III | - | 1 (10%) | - |

Data are presented as mean ± SD or n (%) as appropriate.

We applied the Intrinsic Ignition framework [22] to measure dynamical complexity, focusing on two key metrics: intrinsic ignition and metastability (see Figure 1 and Section 4.4). These metrics quantify (1) a region’s ability to propagate activity and (2) the dynamic balance between segregation and integration in brain networks. Intrinsic Ignition evaluates how a brain region can propagate activity to drive global integration, highlighting its role in shaping hierarchical and dynamic communication. Metastability assesses the temporal variability in global synchronization, reflecting the balance between stable coordination and flexible transitions, which supports adaptive and hierarchical information processing. Together, these metrics offer key insights into the brain’s dynamic complexity.

**Figure 1.**
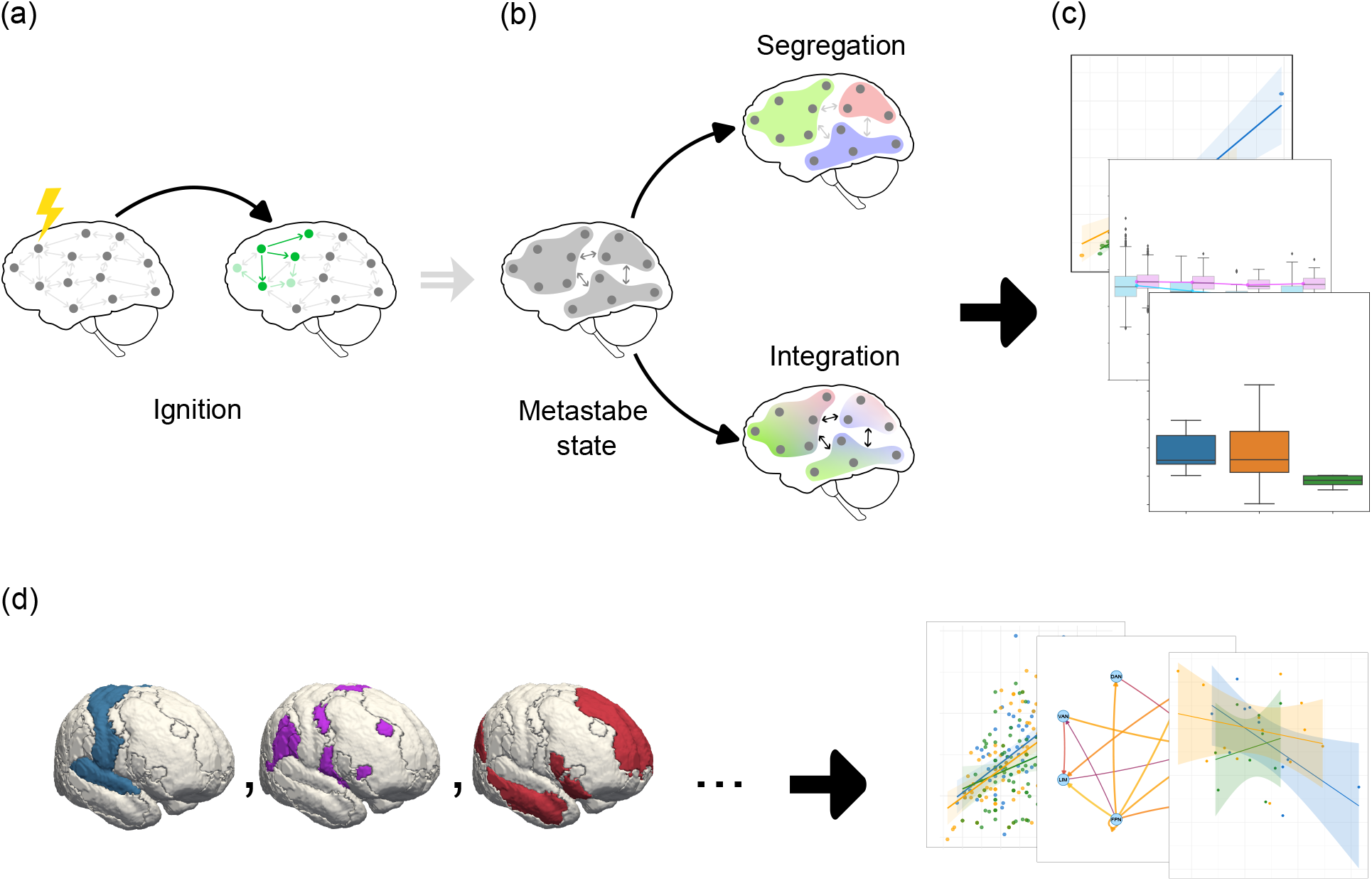
Conceptual illustration of the Intrinsic Ignition Framework and an overall study design. **(a) Intrinsic Ignition**: This metric measures a node’s capacity to propagate information within the network. Initially, nodes are in a basal state, receiving baseline signals. Upon reaching an excitation threshold (indicated by the yellow symbol), the node generates a significant activity spike (ignition), propagating this activity to neighboring nodes over a defined time window. Subsequent nodes may continue to process and propagate this activity. **(b) Metastability**: This metric captures the temporal variability in global synchronization within brain networks, reflecting the dynamic balance between periods of stable synchronization and transitions to more flexible states. It supports both segregation (specialized intra-module processing) and integration (enhanced cross-module communication), while also being linked to the hierarchical organization of brain networks. The illustration depicts brain nodes in a metastable state, transitioning between segregation and integration, highlighting the variability of network dynamics and the adaptive flexibility of the system. **(c) Whole-brain dynamics**: This involves the application of intrinsic ignition (a) and metastability (b) to perform whole-brain analysis at both the subject and regional levels, comparing three groups: control, meningioma, and glioma patients. **(d) Resting State Networks**: Analysis of brain dynamics is conducted using these metrics within specific resting-state networks.

### 2.2 Whole-brain dynamics at the subject level

We assessed the intrinsic ignition and metastability to study whole-brain dynamics differences among the control, glioma, and meningioma groups. Comparing the mean intrinsic ignition by subject among the three groups, as shown in Figure 2a, revealed a pronounced reduction in mean intrinsic ignition in the glioma group compared to controls (*p* = 0.0073), indicating a substantial disruption in whole-brain dynamics. Additionally, a significant decrease in mean intrinsic ignition was noted between meningioma and glioma patients (*p* = 0.0108), but not between meningioma and control groups (*p* = 0.7035), suggesting relatively preserved functional dynamics in meningioma patients. These results are consistent with the computed effect sizes from the sensitivity analysis (see Section 4.5). Between the meningioma and control groups, with a mean variability of 0.024, a minimum detectable difference of 0.023 is required, but only a difference of 0.0034 was obtained. Conversely, for the glioma and control groups, with a mean variability of 0.017, a minimum detectable difference of 0.018 is needed, and a difference of 0.022 was obtained, aligning with the sensitivity analysis.

**Figure 2.**
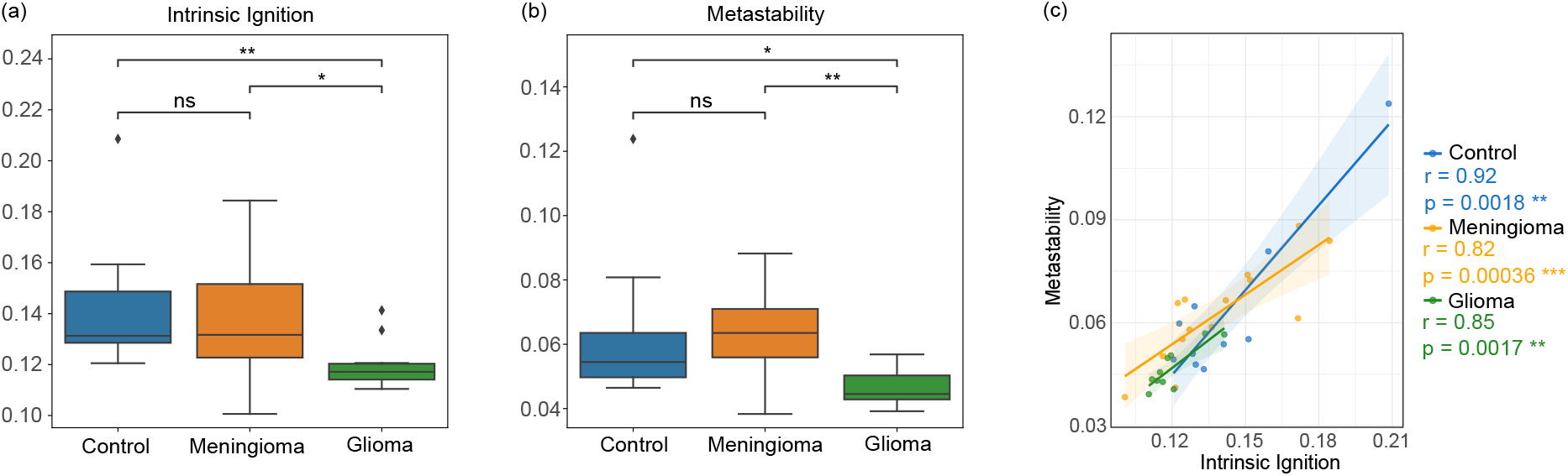
**(a)** Mean intrinsic ignition by subject; **(b)** Mean metastability by subject, analyzed across three groups: control, meningioma, and glioma. Subjects with glioma are statistically distinguishable based on both metrics. **(c)** Correlation among the three groups at the subject level between mean intrinsic ignition and mean metastability, showing a strong correlation in all groups. The legend for the p-values is as follows: *ns* indicates *p* ≥ 0.05, a single asterisk (*) represents 0.01 *< p* ≤ 0.05, two asterisks (**) denote 0.001 *< p* ≤ 0.01, and three asterisks (***) signify *p* ≤ 0.001.

Similarly, the comparison of mean metastability at the subject level between the three groups, shown in Figure 2b, revealed no significant difference between the meningioma and control groups (*p* = 0.3641). However, significant differences were observed between the meningioma and glioma groups (*p* = 0.0065) and between the control and glioma groups (*p* = 0.0173). These results suggest that glioma patients exhibit altered global metastability compared to control and meningioma groups, while meningioma patients maintain metastability levels comparable to those of healthy controls. According to the sensitivity analysis, with a mean variability between the meningioma and control groups of 0.018, a detectable difference of 0.017 is needed, but only a difference of 0.00038 was obtained. Conversely, between the glioma and control groups, with a mean variability of 0.014, a needed difference of 0.015 is required, and a difference of 0.016 was obtained.

Subsequently, we explored the relationship between intrinsic ignition and metastability at the subject level. A Pearson correlation analysis between these two measures, shown in Figure 2c, uncovered a significant positive relationship across the three distinct groups: control (r = 0.92, p = 0.0018), meningioma (r = 0.82, p = 3.588e-04), and glioma (r = 0.85, p = 0.0017). Despite a slight influence of tumor presence on the correlation for both meningiomas and gliomas, all three groups exhibited a robust correlation. However, this scenario changes remarkably when the same correlation analysis is applied at the regional level within the resting-state network for glioma subjects, analyzed in Section 2.4, where a weak correlation is detected. Although a part of this effect can be attributed to the difference in the brain atlas used, this result may also suggest that averaging across all nodes may dilute the impact, leading to an overall strong correlation between these measures.

### 2.3 Whole-brain dynamics at the region level

Next, we investigated the influence of tumor affectation on whole-brain dynamics. The region-level analysis comprised two stages. Initially, we aimed to discern any visible trends in tumor affectation. Regions were classified into four categories based on the extent of tumor involvement: regions without tumor involvement [0.0], regions with up to one-third involvement (0.0, 0.33), regions with one-third to two-thirds involvement [0.33, 0.66), and regions with two-thirds to complete involvement [0.66, 1.0] (see Section 4.4). This categorization was independently applied to both the meningioma and glioma groups.

Figure 3a illustrates the changes for the meningioma group, with delta intrinsic ignition depicted in blue and delta metastability in pink. Here, the effects are negligible (close to zero) until the tumor involvement exceeds one-third for intrinsic ignition and two-thirds for metastability, indicating a more pronounced impact in intrinsic ignition. Conversely, Figure 3b presents the data for the glioma group. Notably, even in regions with no direct tumor affectation, the impact on the metrics is evident, with both showing values below 0.0 across all bins. This pattern suggests a relatively consistent effect across varying degrees of tumor affectation, potentially reflecting the infiltrative nature of gliomas compared to meningiomas, without a significant decline at the higher end of tumor involvement.

**Figure 3.**
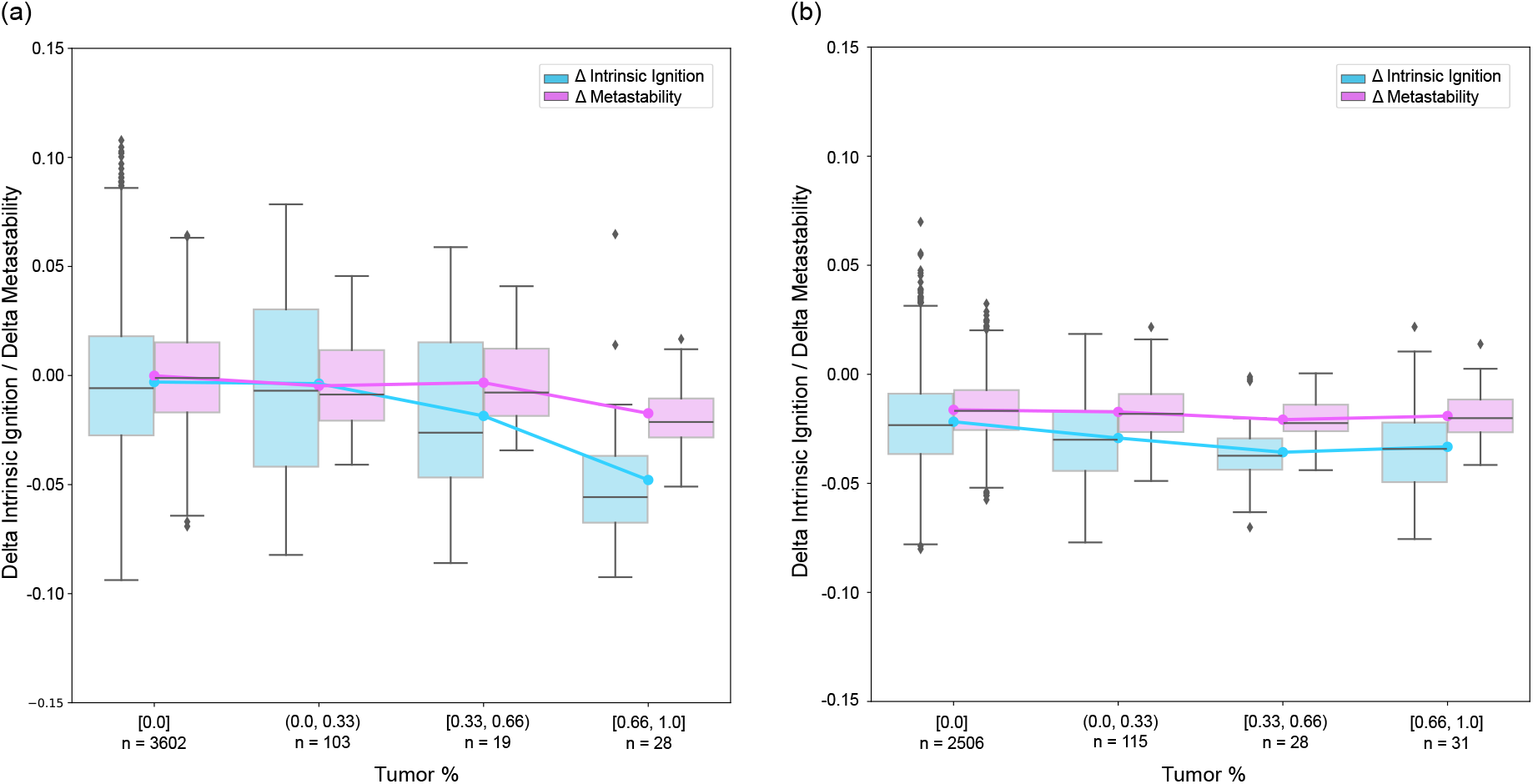
The figure depicts how the mean of the delta intrinsic ignition and delta metastability respond at a regional level as tumor percentage within a region increases. Tumor percentage has been categorized into four bins: [0.0], (0.0, 0.33), [0.33, 0.66), and [0.66, 1.0]. The *n* label on the x-axis represents the number of regions in each bin. Panel (a) illustrates the meningioma group, showing that for a tumor affectation of up to a third, there is little difference from the control group (close to 0.0). The effects on intrinsic ignition and metastability become more pronounced for higher tumor affectations. Panel (b) features the glioma group, where there is a noticeable reduction in both delta intrinsic ignition and metastability compared to the control group, even in regions without tumor affectation, indicating an overall impact on this group. Contrary to the meningioma group, tumor affectation in the glioma group only shows some effect on the delta intrinsic ignition.

To gain a detailed understanding of the impact of tumor percentage on the two neurological response variables, delta intrinsic ignition, and delta metastability, we employed linear mixed-effects model analyses as outlined in Section 4.5 for both meningioma and glioma patient groups. In the meningioma group, there was a statistically significant decrease in delta intrinsic ignition associated with increased tumor percentage, whereas no significant effect was observed for delta metastability. Conversely, neither response variable demonstrated a statistically significant change with varying tumor percentages in the glioma group. However, significant intercepts indicated a baseline shift in both delta measures, independent of tumor percentage. These findings are corroborated by Figure 3, which visually supports the statistical analysis. The subsequent sections provide a detailed description of these results.

#### Meningioma Group

Delta intrinsic ignition demonstrated a significant negative association with tumor percentage (Estimate = -0.03246, SE = 0.01209, t = -2.686, p = 0.035), suggesting that an increase in tumor percentage correlates with a decrease in intrinsic ignition, where each percentage point increase in tumor size reduces this value by approximately 0.032 units. Regarding random effects, significant variability was noted between subjects and across regions, with standard deviations of 0.02481 and 0.00695, respectively. The greater variability at the subject level indicates that individual differences significantly affect the observed ignition values. Additionally, a negative correlation (−0.39) between the random slope of tumor percentage and the intercept at the subject level implies that subjects with higher average tumor percentages generally have lower baseline intrinsic ignition values. The residuals, ranging from -3.99 to 3.37, were mostly close to zero, indicating a good model fit. As shown in the diagnostic plots (Figure 4), the residuals were symmetrical around the median, supporting the validity of the findings that tumor percentage significantly affects Δ intrinsic ignition.

**Figure 4.**
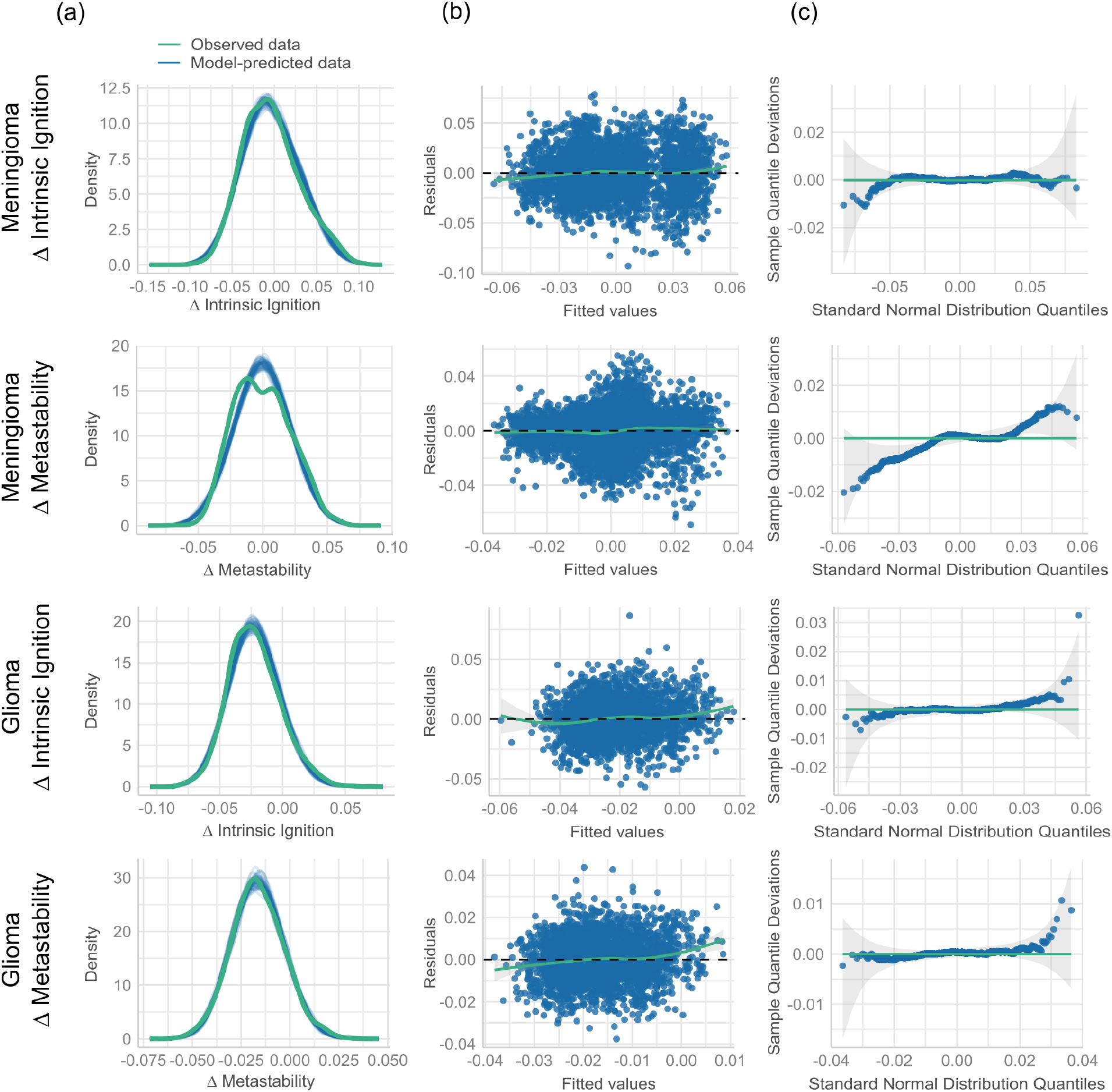
Diagnostic plots for the linear mixed-effects models applied to delta intrinsic ignition and delta metastability. **(a)** The first column illustrates the density plots of observed versus model-predicted values, demonstrating overall alignment. **(b)** The second column presents residuals plotted against fitted values, where deviations from linearity are most evident for glioma - Δ metastability, suggesting a potential violation of the linearity assumption in this case. **(c)** The third column features quantile-quantile (Q-Q) plots of residuals, where the meningioma - Δ metastability model demonstrates some departures from normality.

For delta metastability within the meningioma group, no significant effects of tumor percentage were observed (Estimate = -0.00503, SE = 0.00379, t = -1.328, p = 0.387), indicating that tumor percentage does not impact this measure as strongly as it affects delta intrinsic ignition. Residuals for delta metastability varied from -4.30 to 3.55. However, as shown in the diagnostic plots (Figure 4c), the quantile-quantile (Q-Q) plot for this model demonstrated deviations from normality, suggesting potential issues with the normality assumption of the residuals. Additionally, the predictive checks for this model indicated discrepancies between the observed and model-predicted values, as highlighted by the density plots (Figure 4a), suggesting that the model does not fully capture the observed variability in delta metastability. Despite these issues, the median residual remained slightly positive, indicating that the model fits the data reasonably well.

#### Glioma Group

In the glioma group, the effect of tumor percentage on delta intrinsic ignition was not statistically significant (Estimate = -0.01189, SE = 0.00858, t = -1.386, p = 0.211), indicating that changes in tumor percentage do not predict changes in intrinsic ignition. However, a significant negative intercept (Estimate = -0.02220, SE = 0.00317, t = -7.011, p = 4.9e-05) suggests a consistently lower baseline level of intrinsic ignition across the sample, irrespective of tumor percentage. The residuals in this group varied more widely, from -3.45 to 5.26, with a median very close to zero. The residuals clustered tightly around the median, indicating that the model adequately predicts delta intrinsic ignition for most of the data, despite some residuals suggesting minor deviations from a perfect fit, as shown in the diagnostic plots (Figure 4b).

Similarly, for delta metastability, the effect of tumor percentage was not significant (Estimate = -0.00362, SE = 0.00429, t = -0.845, p = 0.433), but with a statistically significant negative intercept (Estimate = -0.01639, SE = 0.002025, t = -8.095, p = 1.38e-05), indicating a reduction in metastability that is consistent across the glioma group. The negative correlation (−0.57) between the tumor percentage effects and the subject intercepts further suggests that individuals with higher tumor percentages tend to exhibit lower baseline metastability levels. As shown in the diagnostic plots (Figure 4), plotting the residuals versus fitted values demonstrates small deviations from linearity, suggesting minor issues with the linearity assumption. Despite this, the residuals are well-distributed, ranging from -3.53 to 4.10, with a median near zero. The clustering of residuals around the median indicates that the model captures the general trends in the data effectively, though the observed deviations highlight areas where the model could perform better.

The predictive checks for the models revealed that the simulated data closely matched the observed data for most models, as shown in the diagnostic plots (Figure 4). For delta intrinsic ignition in both meningioma and glioma groups, the density plots indicated good alignment between the observed and model-predicted values, confirming the models’ ability to capture the overall trends in the data. Similarly, for delta metastability in the glioma group, the predictive checks supported the adequacy of the model, with a good match between observed and predicted values. However, for delta metastability in the meningioma group, predictive checks revealed some discrepancies, indicating that the model struggled to fully replicate the observed data distribution.

### 2.4 Resting State Networks

A subsequent phase of our analysis utilized Resting State Networks (RSNs), as depicted in Figure 5b and elaborated in Section 4.5. This approach yielded further insights beyond those obtained from the previous analysis described in Section 2.3. Figure 5c illustrates the distribution of tumors at the RSN level: the left panel displays the number of subjects with each specific RSN affected. In contrast, the right panel shows the frequency distribution of the affected RSNs per subject. Figure 5d presents the distributions of the two measures, intrinsic ignition and metastability, across multiple resting-state networks for control, meningioma, and glioma groups. Each data point represents an individual participant, with median values indicated by black bars. The figure illustrates the variability within and across groups, providing an overview of how whole-brain complexity indices are distributed among different clinical populations.

**Figure 5.**
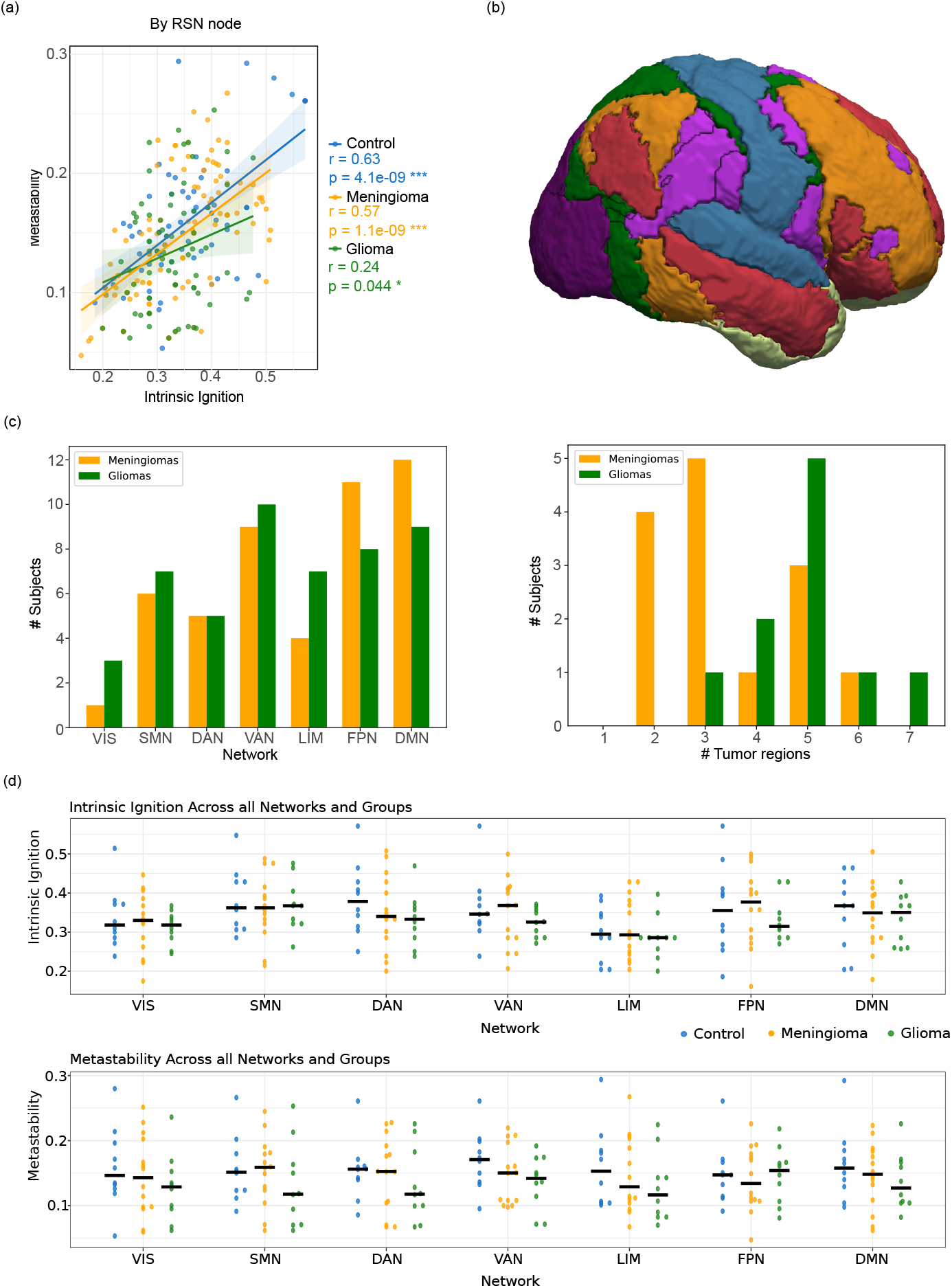
**(a)** Correlation among the three groups at the node level using the Yeo7 parcellation. Correlations are disrupted compared to the control group, especially for glioma. **(b)** Side view of the Yeo7 parcellation: dark purple represents the Visual network, blue the Somatomotor network, dark green the Dorsal attention network, light purple the Ventral attention network, light green the Limbic network, orange the Frontoparietal network, and red the Default Network network. **(c) Left:** This graph presents the number of subjects with a tumor in each network, as indicated on the x-axis. **Right:** This graph displays the number of affected network per subject, with the number of regions indicated on the x-axis. **(d)** Distribution of whole-brain dynamical complexity measures across each resting-state network, with data grouped by subject type (control, meningioma, and glioma). **Top:** Illustrates intrinsic ignition values. **Bottom:** Shows metastability values. Each point represents an individual subject, and median values are indicated by black bars.

Pearson correlation was conducted at the RSN region level, as shown in Figure 5a. Here, a notable difference was observed in subjects with glioma compared to the subject-level analysis in Section 2.2. The control and meningioma groups exhibited moderate correlations, with values of (r = 0.63, p = 4.1e-09) and (r = 0.57, p = 1.1e-09), respectively, suggesting a minimal effect of meningiomas on this correlation. In contrast, the association within the glioma group was significantly weaker (r = 0.24, p = 0.044). This discrepancy likely stems from the divergent effects of gliomas on the two metrics under investigation. Similar outcomes were observed when conducting this analysis with the Shen268 parcellation, indicating that while the subject-level analysis did not show such a weak correlation for the glioma group, it is at the region level.

#### Correlation analysis

A correlation analysis was conducted to explore this phenomenon further and visualize global trends. Graphs representing three distinct groups—control, meningioma, and glioma—were generated. For each group, we examined correlations within and between the following metrics: intrinsic ignition, metastability, and their interaction, as illustrated in Figure 6. In Figure 6a, the inter-region correlation of intrinsic ignition was measured. Although no significant patterns were observed, some weaker correlations were noted in the glioma group, suggesting that inter-region ignition correlations at the global level are generally weak and have high variability among subjects. Higher ignition values in some regions did not correlate with higher values in others.

**Figure 6.**
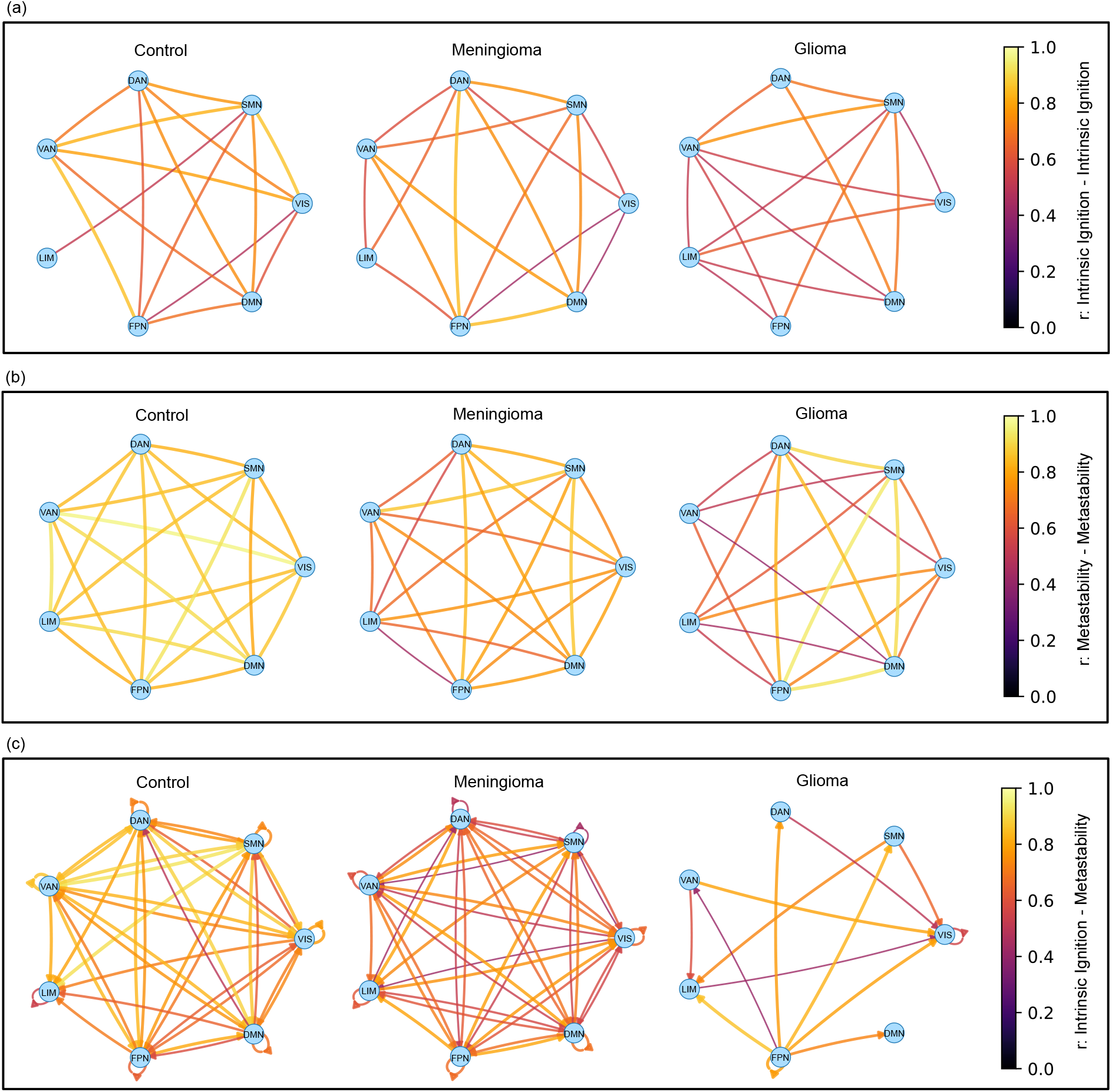
Correlation analyses of intrinsic ignition and metastability, where regions are represented as nodes and correlation strengths as edges, across three distinct groups: control, meningioma, and glioma. **(a)** Inter-region correlations of intrinsic ignition demonstrate generally weak patterns, with minimal notable exceptions. **(b)** Inter-region metastability correlations show strong, consistent correlations in the control group, indicating preserved metastability; these correlations are noticeably weaker in the meningioma group and significantly disrupted in the glioma group. **(c)** Intra- and inter-region correlations between intrinsic ignition and metastability reveal strong correlations in the control group; in contrast, the meningioma group exhibits weaker correlations, and the glioma group displays predominantly disrupted correlations. These visualizations underscore substantial differences in neural network dynamics among the groups.

Inter-region metastability correlation, however, presents a contrasting narrative. As shown in Figure 6b, a strong correlation among all nodes is depicted for the control group, indicating that global levels of metastability are conserved in healthy subjects. In contrast, although the meningioma group also shows correlations between all regions (with values ≥4), these correlations are visually weaker than in the control group, indicating a diminished relationship in this group. Similarly, the glioma group exhibits disrupted correlations among some nodes and generally weaker correlations than the control group.

Finally, Figure 6c visualizes the intra- and inter-region correlations between intrinsic ignition and metastability. The results show similarities to the previous metastability visualization but with more pronounced differences between groups. The control group displays a highly correlated graph between these two measures, suggesting that similar levels in other regions generally mirror regions with higher ignition or metastability. The meningioma group, although still showing these correlations, exhibits weaker connections. Conversely, these correlations are mostly disrupted in the glioma group, marking a significant deviation from the other two groups.

#### CANTAB RVP Mean Latency Test vs Intrinsic ignition

Finally, we investigated the potential association between ignition and cognitive performance using the CANTB Rapid Visual Information Processing (RVP) test, specifically the mean time response for each subject was used. The RVP test assesses sustained attention and information processing speed by requiring participants to monitor a sequence of digits displayed at a rate of 100 digits per minute and respond to specific target sequences (e.g., 2-5-7, 4-3-8) by pressing a button. This test is particularly effective for engaging key brain networks involved in these cognitive functions and has been utilized in previous disease studies [23, 24]. Our analysis focused on the involvement of specific RSN during this task: the Visual network, crucial for processing visual stimuli; the Dorsal network, key in directing and maintaining attention [25]; and the Default Mode Network, often referred to as the “task-negative” network due to its typically reduced activity during focused attention tasks [26, 27]. Notably, two subjects—one diagnosed with a meningioma (age: 49, tumor size: 0.75 cm^3^, female) and another with a glioma (age: 53, tumor size: 15.16 cm^3^, male)—were unable to complete the test. As illustrated in Figure 7, a significant negative correlation was observed between intrinsic ignition and RSN activity in the control group in the key regions: the Visual (r = -0.67, p = 0.0328; see Figure 7a), the Dorsal region (r = -0.70, p = 0.0234; see Figure 7b), and the Default Network (r = -0.76, p = 0.0103; see Figure 7c). This association was not observed in the groups with meningiomas and gliomas, suggesting a potential differential impact of these conditions on cognitive network functionality. Analysis of other networks did not show any significant correlations.

**Figure 7.**
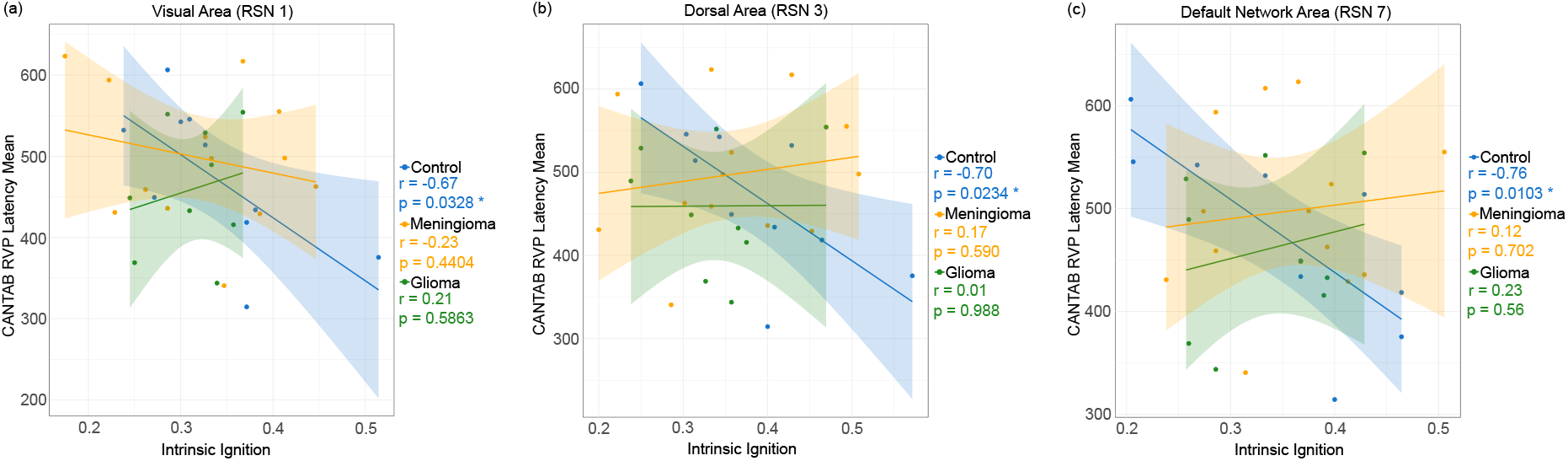
This graph illustrates the correlation between the RVP Latency Mean test scores and ignition across regions of interest within the Resting State Network (RSN) using the Yeo7 parcellation. The control group demonstrates a strong negative correlation, whereas this correlation is notably disrupted in the meningioma and glioma groups. **(a)** Visual Network (RSN 1) **(b)** Dorsal Network (RSN 3) **(c)** Default Mode Network (RSN 7)

## 3 Discussion

In this work, we studied the dynamical complexity underlying the whole-brain functional network to understand the impacts of glioma and meningioma brain tumors on brain information processing. We comprehensively analyzed the subject, region, and resting-state network levels. Our findings reveal significant differences in how these tumors disrupt brain function, highlighting the distinct pathological mechanisms of gliomas and meningiomas. At the subject level, glioma patients exhibited significant reductions in intrinsic ignition and metastability compared to controls, indicating widespread disruptions in brain dynamics. Meningioma patients, in contrast, displayed relatively preserved metrics, with no significant differences from controls, suggesting a more localized impact. At the region level, tumor burden was associated with reduced intrinsic ignition in meningioma patients, particularly in regions with over one-third involvement. Glioma patients exhibited widespread reductions in both metrics, even in tumor-free regions, reflecting their diffuse and invasive nature. At the RSN level, glioma patients displayed severely disrupted metastability correlations, while meningioma patients exhibited milder disruptions. Correlations between intrinsic ignition and metastability, robust in controls, were weakened in both patient groups, particularly in gliomas.

Our subject-level analysis revealed pronounced differences in brain dynamics among control, meningioma, and glioma groups. Subjects with gliomas exhibited significantly reduced intrinsic ignition and metastability compared to controls. This suggests a widespread disruption of brain network dynamics, reflecting the glioma’s aggressive and infiltrative nature. These findings align with clinical observations that gliomas, due to their diffuse growth patterns, often lead to extensive neurological impairments [28]. In contrast, meningioma subjects did not show significant differences from controls in these metrics, indicating that meningiomas, typically more localized, have a less pervasive impact on global brain function. This supports the clinical perspective that meningiomas cause focal symptoms depending on their location and size [29]. Our results are consistent with previous research on neural network alterations underlying cognitive deficits in brain tumor patients [30].

At the regional level, the extent of tumor involvement differentially affected ignition and metastability. In meningioma patients, significant disruptions in intrinsic ignition were observed in regions with more than one-third of tumor involvement. Metastability was affected only in regions with the highest tumor burden, suggesting that meningiomas maintain relatively normal network dynamics until a critical threshold is reached. This localized impact aligns with the clinical behavior of meningiomas [29], which tend to be more contained. Conversely, glioma patients showed widespread reductions in both metrics, even in regions without direct visible tumor involvement as previously demonstrated [31, 32]. Excessive peritumoral glutamate release by gliomas [33, 34] could also add to this effect. This widespread impact reflects the invasive nature of gliomas, which disrupt brain function through various mechanisms including diffuse infiltration and alteration of surrounding neural tissue. Our linear mixed-effects model analyses supported these observations, showing significant associations between tumor burden and intrinsic ignition reduction in meningioma patients. In glioma patients, consistent disruptions were observed regardless of tumor percentage, underscoring the extensive network perturbations caused by these tumors. Similarly, Aerts et al. [35] demonstrated that individual biophysical model parameters reveal distinct focal and distant effects on local inhibitory connection weights in tumor-affected brains.

The RSN-level analysis provided further insights into the differential impacts of gliomas and meningiomas on brain network dynamics. In the control group, strong and consistent correlations in metastability across all RSN nodes indicated preserved functional connectivity and network stability. The meningioma group showed slightly weaker correlations, suggesting localized disruptions without extensive network-wide impacts. However, the glioma group exhibited significantly disrupted correlations, underscoring the extensive network perturbations induced by these tumors.

Furthermore, correlation analysis between intrinsic ignition and metastability further emphasized these differences. The control group demonstrated strong positive correlations, reflecting integrated and stable network dynamics. In contrast, the glioma group showed disrupted correlations, indicating a decoupling of network dynamics and reduced stability. The meningioma group exhibited intermediate patterns, consistent with their more localized impact on brain function. These findings suggest that gliomas fundamentally alter the brain’s intrinsic network architecture, leading to widespread functional connectivity and stability disruption. It should be noted that part of these disrupted correlations could also result from brain compensatory mechanisms, as hypothesized by Duffau et al. [36].

Finally, our investigation into the relationship between intrinsic ignition and cognitive performance, as measured by the CANTAB RVP test, revealed significant negative correlations in the control group across key RSN regions, including the Visual, Dorsal Attention, and Default Mode networks. This suggests that higher ignition is associated with better cognitive performance in healthy individuals. However, this relationship was disrupted in both tumor groups, indicating that the presence of tumors affects the functional integration of cognitive networks. The absence of significant correlations in the meningioma and glioma groups suggests that these tumors impair the brain’s ability to maintain stable and integrated network dynamics, which is essential for optimal cognitive function. These findings highlight the potential of intrinsic ignition as a biomarker for cognitive deficits in brain tumor patients, providing a quantitative measure of the functional impact of tumors on cognitive networks.

This study is not without limitations. First, the choice of brain atlases, such as Shen268 and Yeo7, may limit the granularity and specificity of the observed effects, potentially overlooking finer-scale dynamics. Second, the sample size is relatively small, particularly for subgroup analyses, which may limit statistical power and the generalizability of these findings. Third, variations in MRI acquisition parameters, particularly the differences in TR values across participants, may introduce inconsistencies in the measured metrics. Addressing these limitations in future studies through larger, more diverse samples and standardized imaging protocols will enhance the robustness and applicability of these findings.

In summary, this study demonstrates that gliomas and meningiomas differentially affect brain dynamics: gliomas cause widespread network disruptions while meningiomas have a more localized impact. These findings have important implications for understanding the pathophysiology of brain tumors and developing targeted therapeutic strategies. Future research should explore the potential of intrinsic ignition and metastability as biomarkers for monitoring disease progression and treatment response in brain tumor patients. Additionally, larger and more diverse cohorts are needed to validate these findings and to explore the effects of different tumor grades and locations on brain network dynamics.

## 4 Materials and Methods

### 4.1 Participants

Data for this study was obtained from a dataset [19] comprising pre-operative information from individuals diagnosed with glioma, meningioma, and control subjects. The control group consisted of partners of the patients to ensure comparable levels of emotional distress. Participants were enrolled at Ghent University Hospital (Belgium) from May 2015 to October 2017. Inclusion criteria required participants to be 18 years or older, diagnosed with a supratentorial meningioma (WHO grade I or II) or glioma (WHO grade II or III) brain tumor, capable of completing neuropsychological assessments, and medically cleared for MRI examination.

For this study, 10 control subjects (mean (SD) age = 59.7 (10.1); 3 females), 10 glioma subjects (mean (SD) age = 47.7 (12.3); 4 females), and 14 meningioma subjects (mean (SD) age = 60.4 (12.3); 11 females) from the dataset [19] were included. Among the glioma subjects, diagnoses included 1 case of Oligo-astrocytoma (WHO grade II), 1 case of Ependymoma (WHO grade II), 3 cases of Anaplastic Astrocytoma (2 with WHO grade III and 1 with WHO grade II-III), 3 cases of Oligodendroglioma (WHO grade II), 1 case of glioma (WHO grade III), and 1 case of Astrocytoma (WHO grade II). Among the meningioma subjects, diagnoses included 13 cases of WHO grade I meningioma and 1 case of WHO grade II meningioma. The average tumor size for glioma patients was 23.43 *cm*^3^ (SD = 18.24), while for meningioma patients, it was 17.01 *cm*^3^ (SD = 29.17) see Table 1.

### 4.2 MRI acquisition

From the above dataset, the following acquired and derivative data were utilized in this study: T1-weighted Magnetization Prepared Rapid Gradient Echo (MPRAGE) anatomical scans (160 slices, TR = 1750 ms, TE = 4.18 ms, field of view = 256 mm, flip angle = 9°, voxel size = 1 × 1 × 1 mm, TA = 4:05 min). Resting-state functional echo-planar imaging (EPI) scans were acquired in interleaved order (42 slices, TE = 27 ms, field of view = 192 mm, flip angle = 90°, voxel size = 3 × 3 × 3 mm). It is noted that, as stated by the original authors, the initial acquisition included 4 control subjects, 5 meningioma subjects, and 2 glioma subjects with a TR of 2100 ms, resulting in a TA of 6:24 min. Subsequent acquisitions were accidentally conducted with a TR of 2400 ms, resulting in a TA of 7:19 min. Tumor masks in MNI spaces (originally obtained through a combination of manual delineation and disconnectome) were also used. More specific information on the dataset can be found in [35, 37, 38].

The dataset includes an assessment of cognitive performance using the Cambridge Neuropsychological Test Automated Battery (CANTAB; Cambridge Cognition, 2017). The CANTAB is a computerized battery designed to evaluate various cognitive abilities such as memory, attention, executive function, and decision-making. Specifically, the Rapid Visual Information Processing (RVP) test from the CANTAB was employed in this study. The RVP test measures several outcome variables including the speed of response, sensitivity to the stimuli, and the probability of false alarms. These metrics have been previously demonstrated to effectively assess cognitive functions in various disorders [23, 24]. Our analysis used the mean response speed as the primary measure of cognitive performance.

### 4.3 Functional MRI processing

fMRI data was initially processed using the *FSL* suite (FMRIB Software Library), specifically employing *FEAT* (FMRI Expert Analysis Tool, Version 6.00). The preprocessing steps included: motion correction using *MCFLIRT* [39], interleaved slice timing correction, brain extraction using BET [40], grand-mean intensity normalization across the entire 4D dataset using a single scaling factor, and high-pass temporal filtering with a cutoff of 100 s. Non-linear registration was performed to align the processed data with the MNI152 space. Subsequently, using the Python *Nilearn* library (version 0.10.2), which relies heavily on the *scikit-learn* package [41], signals were z-score standardized and extracted for each of the defined parcellation regions. Two atlases were employed: the Shen268 [42], consisting of 268 regions derived from a group-wise whole-brain graph-theory-based parcellation [20], and the Yeo7 using the liberal version of the atlas [43], comprising seven networks based on resting-state networks [21]. These networks include the Visual Network (VIS, region 1) for visual processing, the Somatomotor Network (SMN, region 2) for bodily movement control, the Dorsal Attention Network (DAN, region 3) for top-down attention, the Ventral Attention Network (VAN, region 4) for bottom-up attention responses, the Limbic Network (LIM, region 5) for emotion and memory, the Frontoparietal Network (FPN, region 6) for complex cognitive functions, and the Default Mode Network (DMN, region 7) for introspective activities during rest.

#### Tumor Regions

The information for each tumor region was extracted using the following criteria: values from the voxel mask in MNI152 space were thresholded to 0.1 and mapped to the atlas parcellation. Regions were identified as tumors if at least one voxel intersection existed, and the affected percentage was calculated at the voxel level.

### 4.4 Dynamical complexity

Dynamical complexity is measured using the Intrinsic Ignition framework [22, 44]. This framework underlies the hierarchical dynamics of neural activity within specific brain states, offering insights into brain network complexity. It assesses the ability of particular brain regions to transmit feedforward and recurrent neural activity, which is pivotal for analyzing the inherent fluctuations and their impacts on the functional network (Figure 1). The framework introduces two critical metrics: Intrinsic ignition and metastability.

Intrinsic Ignition measures the capacity of a brain region to initiate and propagate activity across the network (Figure 1a). This is quantified by assessing how an intrinsic event in a region drives integration across other regions over time, highlighting its role in local-global communication. Regions with higher intrinsic ignition are pivotal for fostering global network integration and are often associated with key nodes in the brain’s structural and functional hierarchy. The hierarchical organization of the brain is reflected in the variability of intrinsic ignition across regions, with regions at higher levels of the hierarchy showing greater capacity for widespread integration.

Metastability, on the other hand, evaluates the temporal variability of global synchronization within the brain network (Figure 1b). It captures the dynamic balance between periods of stable synchronization and flexible transitions between states. High metastability indicates the brain’s ability to dynamically coordinate specialized processes (segregation) while enabling global integration of information across modules. Moreover, metastability reflects the hierarchical organization of the brain, where regions operate at varying levels of complexity and contribute to adaptive and efficient information processing.

This framework has been effectively applied to explain variations in whole-brain dynamics across diverse contexts, such as meditation [45], deep sleep [22], disorders of consciousness [46], and in preterm newborns [47].

Utilizing this framework, we can measure the intrinsic dynamics of brain activity using resting-state fMRI data from control, glioma, and meningioma tumor subjects.

The following steps were performed. First, a Butterworth of order 2 bandpass filter ranging from 0.01 Hz to 0.07 Hz was applied to the time series of each node. This range was chosen to focus on the frequencies relevant to the dynamics of interest in the brain, aligning with previous studies [48]. Still, it is worth noticing that the whole procedure is sensitive to these frequencies. Additionally, strong artifacts were removed, defined as values exceeding three times the standard deviation or falling below minus three times the standard deviation. After, a binarization of the filtered time series was applied. For this, a transformation into z-scores is applied *z*_*i*_(*t*) imposing a threshold *θ* such that *σ*_*i*_(*t*) = 1 if *z*_*i*_(*t*) *> θ* and is crossing the threshold from below, *σ*_*i*_(*t*) = 0 otherwise [49]. Using previous event binarization, a connection matrix was constructed for each time frame to compute integration. The ignition for each region was calculated as the mean of the maximum integration within a specific time window (*tw*) for each event.

As before, the metastability associated with a region was quantified as the standard deviation of the integration. To ensure that all subjects share the same time window duration despite differences in fMRI TRs used during data acquisition, we chose a *tw* such that it is 7 for subjects with a TR of 2400 ms and 8 for those with a TR of 2100 ms. This ensures standardization across all subjects in terms of time window duration.

To further explore the relationship between intrinsic ignition, metastability, and tumor involvement, we conducted a detailed analysis at the level of individual brain regions, building on findings from the subject-level analysis. Our objective was to visualize the impact of tumor size on the metrics of each region. Given the inherent variability among different brain regions, we standardized intrinsic ignition and metastability values based on their changes relative to the control group. For each of the 268 regions, the change was calculated as 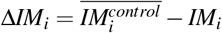, where *IM* represents either *intrinsic ignition* or *metastability*, and *i* denotes the region in the parcellation. The control mean of a region, 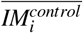, is derived from the control group.

To elucidate the trends in tumor involvement, we categorized the regions into four bins based on the extent of tumor affectation: regions without tumor [0.0], regions with up to one-third tumor involvement (0.0, 0.33), regions with one-third to two-thirds involvement [0.33, 0.66), and regions with two-thirds to complete involvement [0.66, 1.0]. We then analyzed how the mean delta values Δ*IM*_*i*_ changed across these categories. This analysis was performed separately for each tumor type: meningiomas and gliomas.

### 4.5 Statistics

We conducted our statistical analyses at three distinct levels: subject-level, region-level, and resting-state network (RSN)-level. At all levels, we considered two key metrics derived from the Intrinsic Ignition Framework: intrinsic ignition and metastability. Although both metrics are generated from the same underlying data, they capture fundamentally different dimensions of brain dynamics. Specifically, intrinsic ignition reflects the mean level of integration elicited by a brain region (or set of regions), whereas metastability quantifies the temporal variability in global synchronization. Because these metrics represent orthogonal properties (mean vs. variability), correlation analyses between intrinsic ignition and metastability are methodologically sound and provide valuable insights into how these different dimensions of brain dynamics relate to each other.

#### Subject-level

In our study, we first investigated the two metrics of the Intrinsic Ignition Framework at the subject level using the Shen268 parcellation, which comprises 268 nodes. This initial phase aimed to identify intrinsic ignition and metastability differences among the control, glioma, and meningioma groups. For this purpose, we employed the Mann-Whitney-Wilcoxon two-sided tests. This non-parametric method was chosen due to its suitability for data not assumed to follow a normal distribution. Using the Benjamini-Hochberg correction technique, we mitigate the risk of Type I errors, which are inherently increased by multiple comparisons.

To ensure a sufficient sample size for our subject-level analysis, we used G*Power [50] to conduct a sensitivity analysis based on the Wilcoxon-Mann-Whitney test (two groups). We set the power to 1 − *β* = 0.8 and the significance level to *α* = 0.05, resulting in minimum effect sizes of *d* = 0.97 for the meningioma group and *d* = 1.06 for the glioma group.

#### Region-level

To conduct the statistical analysis for region-level based analysis, we considered the potential risk of statistical pseudoreplication due to collecting multiple data points from each participant, which are not independent [51]. To address this issue, we utilized linear mixed-effects models (LMMs) implemented in the R software package. These models effectively manage the data’s hierarchical structure by incorporating fixed effects, such as differences between groups, and random effects, which account for variability among subjects.

The models were estimated using restricted maximum likelihood (REML) methods, and t-tests were performed using Satterthwaite’s method to approximate degrees of freedom. In terms of model-predicted values, both absolute and delta values were considered. Given slightly better predictive outcomes with the delta values, these were used for further analyses. When comparing multiple models through ANOVA, it became evident that while delta values (Δ*IM*_*i*_) captured some variability among regions, incorporating the region ID into the model enhanced its predictive power. This suggests that delta values alone are insufficient to account for all variability. For both groups (meningioma and glioma), we conducted the analysis separately to accommodate potential variances in tumor biology and its impacts on brain dynamics.

The final specification of our linear mixed-effects models was as follows:

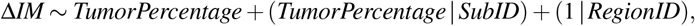

where *IM* represents either intrinsic ignition or metastability as explained in Section 4.4, *TumorPercentage* denotes the percentage of tumor involvement for a region, *SubID* is the subject’s ID, and *RegionID* is the ID of the region.

#### Resting State Networks

To visualize the global trends of dynamical complexity (intrinsic ignition and metastability) across different brain networks, we utilized the Yeo7 parcellation [21], which segments the brain into 7 networks as explained in Section 4.3. This offered two key advantages: firstly, its consolidation into only 7 regions allows for a more focused examination of the specific impacts of tumors, albeit at the expense of granularity; secondly, this parcellation corresponds to well-defined resting-state networks (RSNs), associating each network with distinct cognitive functions such as visual, somatomotor, dorsal attention, ventral attention, limbic, frontoparietal, and default mode network, as shown in Figure 5b.

We constructed correlation graphs after an initial general overview using Pearson correlation analysis of the two metrics at the network level for each group. These graphs were generated for three distinct groups: control, meningioma, and glioma. For each group, we examined Pearson correlations within and between the following metrics: intrinsic ignition, metastability, and their interaction. These graphs represent networks as nodes, with edges indicating the correlation’s strength. Edges were included only if the correlation coefficient was moderate or higher (≥0.4). The color and thickness of each edge are directly proportional to the correlation strength, highlighting stronger relationships. All depicted correlations are positive.

## Data availability

This study utilized the Brain Tumor Connectomics Data, which contains pre-operative data from 11 glioma patients, 14 meningioma patients, and 11 control subjects previously described in [35, 37].

The dataset is publicly available at https://doi.org/10.18112/openneuro.ds001226.v5.0.0.

## Code availability

Upon acceptance, all code for implementing and reproducing our results will be available at https://github.com/ajunca/BrainTumor

